# Mutation detection in thousands of acute myeloid leukemia cells using single cell RNA-sequencing

**DOI:** 10.1101/434746

**Authors:** Allegra A. Petti, Stephen R. Williams, Christopher A. Miller, Ian T. Fiddes, Sridhar N. Srivatsan, David Y. Chen, Catrina C. Fronick, Robert S. Fulton, Deanna M. Church, Timothy J. Ley

## Abstract

Virtually all tumors are genetically heterogeneous, containing subclonal populations of cells that are defined by distinct mutations^1^. Subclones can have unique phenotypes that influence disease progression^2^, but these phenotypes are difficult to characterize: subclones usually cannot be physically purified, and bulk gene expression measurements obscure interclonal differences. Single-cell RNA-sequencing has revealed transcriptional heterogeneity within a variety of tumor types, but it is unclear how this expression heterogeneity relates to subclonal genetic events – for example, whether particular expression clusters correspond to mutationally defined subclones^3,4,5,6-9^. To address this question, we developed an approach that integrates enhanced whole genome sequencing (eWGS) with the 10x Genomics Chromium Single Cell 5’ Gene Expression workflow (scRNA-seq) to directly link expressed mutations with transcriptional profiles at single cell resolution. Using bone marrow samples from five cases of primary human Acute Myeloid Leukemia (AML), we generated WGS and scRNA-seq data for each case. Duplicate single cell libraries representing a median of 20,474 cells per case were generated from the bone marrow of each patient. Although the libraries were 5’ biased, we detected expressed mutations in cDNAs at distances up to 10 kbp from the 5’ ends of well-expressed genes, allowing us to identify hundreds to thousands of cells with AML-specific somatic mutations in every case. This data made it possible to distinguish AML cells (including normal-karyotype AML cells) from surrounding normal cells, to study tumor differentiation and intratumoral expression heterogeneity, to identify expression signatures associated with subclonal mutations, and to find cell surface markers that could be used to purify subclones for further study. The data also revealed transcriptional heterogeneity that occurred independently of subclonal mutations, suggesting that additional factors drive epigenetic heterogeneity. This integrative approach for connecting genotype to phenotype in AML cells is broadly applicable for analysis of any sample that is phenotypically and genetically heterogeneous.

---

Four cases of *de novo* AML and one of secondary AML were selected for study (clinical details in https://github.com/genome/scrna_mutations). eWGS (Fig. 1a) showed that these cases were genetically representative of AML, containing on average 26 mutations in well-established driver genes (e.g. *DNMT3A*, *FLT3*, *NPM1*, *TP53, NRAS, IDH1, CEBPA)*, and representative clonal architecture, with at least one detectable subclone per case (Table 1, Supplementary Table 1) ^10^. Bulk RNA-sequencing revealed that on average, fewer than half of the mutations detected by eWGS were expressed (Table 1).

**Figure 1.**
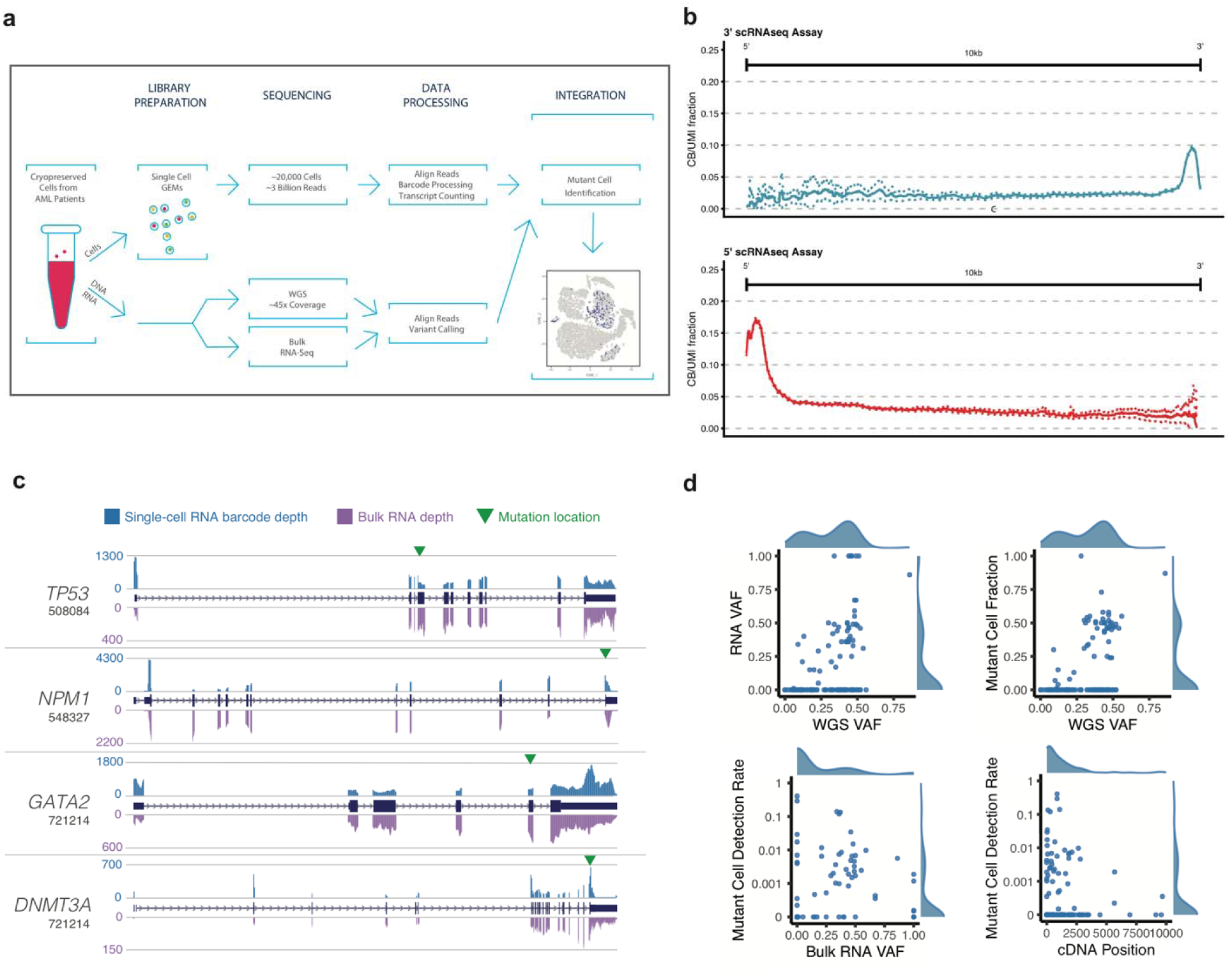
Workflow, coverage, and performance metrics for variant detection in single cells. (a) Cryopreserved bone marrow cells from AML patients underwent eWGS, bulk RNA-seq, and scRNA-seq. Somatic mutations were defined using eWGS data, identified in individual cells using scRNA-seq data, and interpreted in the context of expression heterogeneity. (b) Fraction of unique transcripts (molecules) whose reads map to any given position up to 10 kbp away from the capture site in both the 5’ and 3’ kits. (c) Comparison of single-cell and bulk RNA-seq coverage data for specific genes of interest. (d) Relationship between RNA and eWGS VAF; dependence of Mutant Cell Fraction on eWGS VAF; dependence of Mutant Cell Detection Rate on bulk RNA VAF, and dependence of Mutant Cell Detection Rate on bulk RNA VAF.

**Table 1.** Overview of mutation discovery and detection in eWGS and scRNA-seq data.

| Sample | 508084 | 548327 | 721214 | 782328 | 809653 | Mean [SD] |
| --- | --- | --- | --- | --- | --- | --- |
| No. Cells | 14,964 | 11,620 | 20,474 | 21,731 | 21,038 | 17,965 [3,979] |
| Reads/Cell | 192,427 | 346,965 | 176,035 | 214,284 | 189,751 | 223,892 [62,745] |
| Reads Mapped Confidently to Genome (%) | 84.9 | 87.2 | 92.2 | 79.8 | 90 | 87 [4.29] |
| Reads Mapped Confidently to Transcriptome (%) | 64.2 | 63.7 | 73 | 61.9 | 68.8 | 66 [4.04] |
| Median Genes detected per Cell | 2,405 | 1,383 | 2,260 | 1,376 | 1,829 | 1,851 [428] |
| Total Genes Detected | 22,645 | 22,503 | 23,376 | 25,389 | 23,102 | 23,403 [1041] |
| WGS variants | 19 | 13 | 41 | 31 | 28 | 26.4 |
| Expressed WGS variants (Bulk) | 10 | 5 | 18 | 7 | 8 | 9.6 |
| scRNA-seq variants | 8 | 7 | 17 | 12 | 8 | 10.4 |
| % expressed WGS variants discovered in scRNA-seq | 80% | 140% | 94% | 170% | 100% | 117% |
| Mutant cells per variant | 13-453 | 1-3012 | 1-3944 | 1-2619 | 1-207 | 3.4-2047 |
| Mutant cells per variant (median) | 32 | 48 | 30 | 111 | 21.5 | 48.5 |
| Key WGS Variants (No. cells with scRNA-seq coverage) | *IKBKB*<sup>V616M</sup> (150)<br>*FLT3-ITD* (707)<br>*NUP98-NDS1* (1) | *IDH1*<sup>R132H</sup> (118)<br>*NPM1*<sup>W288fs</sup> (5591)<br>*SRSF2*<sup>P95H</sup> (2349) | *DNMT3A*<sup>R882H</sup> (409)<br>*FLT3-ITD* (479)<br>*FLT3*<sup>F612L</sup> (306)<br>*NPM1*<sup>W288fs</sup> (11,672)<br>*GATA2*<sup>R361C</sup> (1629) | *NRAS*<sup>G12S</sup> (949)<br>*NRAS*<sup>G12D</sup> (951)<br>*U2AF1*<sup>S34F</sup> (4509) | *NRAS*<sup>G12D</sup> (1412)<br>*TP53*<sup>E286G</sup> (239)<br>*CEBPA*<sup>R142fs</sup> (84) | N/A |
| Additional variants with expression signature (number of cells with coverage) | *RNF10* (103) |  |  | *NAGLU*<sup>E634K</sup> (216) |  | N/A |

High-depth sequencing of duplicate scRNA-seq libraries (Table 1) generated using the 5’ (v1) and 3’ (v2) 10x Genomics Chromium Single Cell Gene Expression workflows yielded consistent low-level coverage at least 10 kbp from the 5’ and 3’ ends of the average transcript (Fig. 1b). For the average gene assayed using the 5’ kit, at least 2.5% of the unique sequenced transcripts mapped to any given base up to 10 kbp away from the 5’ transcription start site of the gene. Moreover, the same transcripts had highly correlated coverage patterns in single cell and bulk RNA-seq data (Fig. 1c).

We identified mutation-containing reads and cells by extracting mutant and wild-type Unique Molecular Indices (UMIs) and cell barcodes corresponding to each variant position in the eWGS data (https://github.com/genome/scrna_mutations). A cell was labeled “mutant” if it contained at least one variant-containing read, and “wild-type” if only wild-type reads were detected. Due to low expression and allele dropout, mutations were not detectable in all AML cells; further, “wild-type” cells may contain undetected heterozygous mutations. Most cells with detected somatic mutations contained one mutation, with one read mapping to the variant position (Supplementary Fig. 1). Single nucleotide variants (SNVs), insertions and deletions (indels, including *FLT3*-ITD and *NPMc*), and one gene fusion (*NUP98-NSD1*) were detectable in the single-cell data. We detected an average of 49 mutant cells per variant (range: 1-3944). Founding clone mutations, subclonal mutations, and putative driver mutations were detectable in dozens to thousands of cells in each case (Table 1, Supplementary Table 1). Previously, somatic variant detection from scRNA-seq data involved full-length cDNAs from small numbers of cells^5,11^. Sensitivity of mutation detection was comparable in single cell and bulk RNA-seq data: on average, a slightly higher fraction of known mutations were detected in the scRNA-seq data (Table 1, Fig. 1d, Supplementary Discussion).

We next sought to interpret the mutation data in the context of expression heterogeneity, which we summarized in each case using principal component analysis (Methods). We observed complex relationships among clusters (such as partially overlapping expression signatures), and multiple sources of heterogeneity in all samples, including variable expression of known hematopoietic cell-type markers (e.g. *CD3D* (T-cells), *CD79A* (B-cells), and *HBA1* (erythrocytes)), cell cycle genes (e.g. *TUBA1B*, *TOP2A*), markers of myeloid differentiation (e.g. *AZU1*, *ELANE*, *MPO*, *PRTN3*), mitochondrial genes, and ribosomal genes (Supplementary Fig. 2-6, Supplementary Table 2). This suggested that the distribution of cell types within the bone marrow samples of AML patients is one major source of expression heterogeneity.

To investigate sample composition in a more granular and unsupervised manner, we identified the nearest hematopoietic lineage of each cell by matching it to the most similar lineage-specific expression profile in the DMAP database^12^ (Fig. 2c, Fig. 3). The inferred sample composition varied widely among subjects, particularly with respect to the fraction of lineage-defined cells (e.g. cells resembling myelomonocytic cells, T-cells, B-cells, and erythrocytes). All samples contained clusters of immature cells, including cells resembling hematopoietic stem cells (HSCs), common myeloid progenitors (CMPs), and megakaryocyte-erythroid progenitors (MEPs), which could represent either immature non-malignant cells or AML cells. It is not possible to define AML cells using gene expression patterns alone, and previous approaches for deconvoluting mixtures of tumor and normal cells are not broadly applicable to AML samples, which usually have few copy number alterations^3,4,8,10,13,14^.

**Figure 2.**
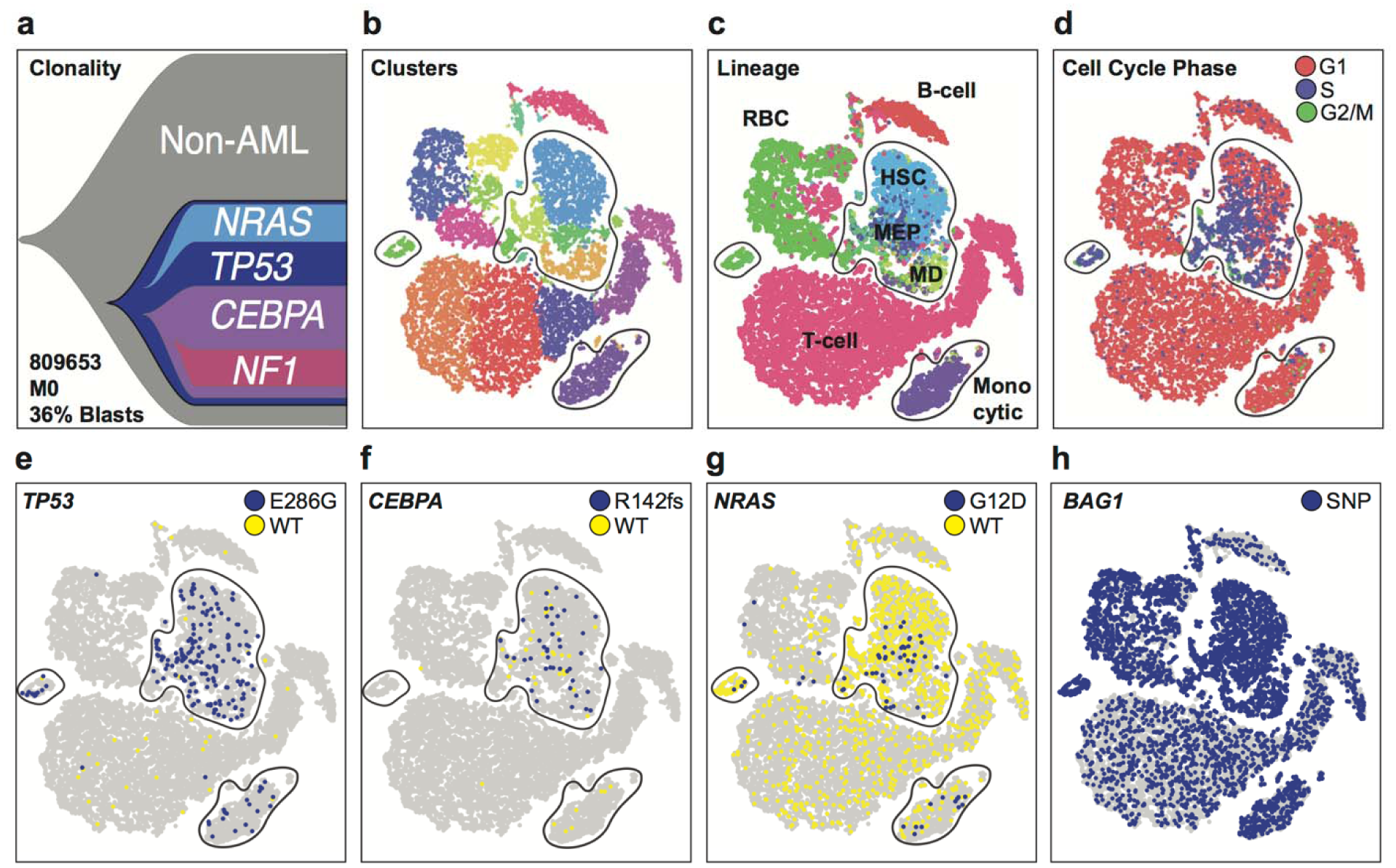
Single-cell mutation detection and interpretation in case 809653. (a) Clonality inferred from eWGS, with driver genes associated with each subclone. (b) t-SNE plot of scRNA-seq data with cells colored according to graph-based cluster assignment. In panels b-g, putative clusters of AML cells are circled. (c) Cells colored according to inferred lineage. (d) Cells colored according to cell cycle phase. (e-g) Cells colored according to single-cell genotype at the *TP53*^E286G^, *CEBPA*^R142fs^, and *NRAS*^G12D^ sites: blue, at least one mutant read detected; yellow, wild-type reads only; gray, no coverage. (h) Cells colored according to single-cell genotype at the homozygous *BAG1* germline SNP: blue, at least one mutant read detected; gray, no coverage.

**Figure 3.**
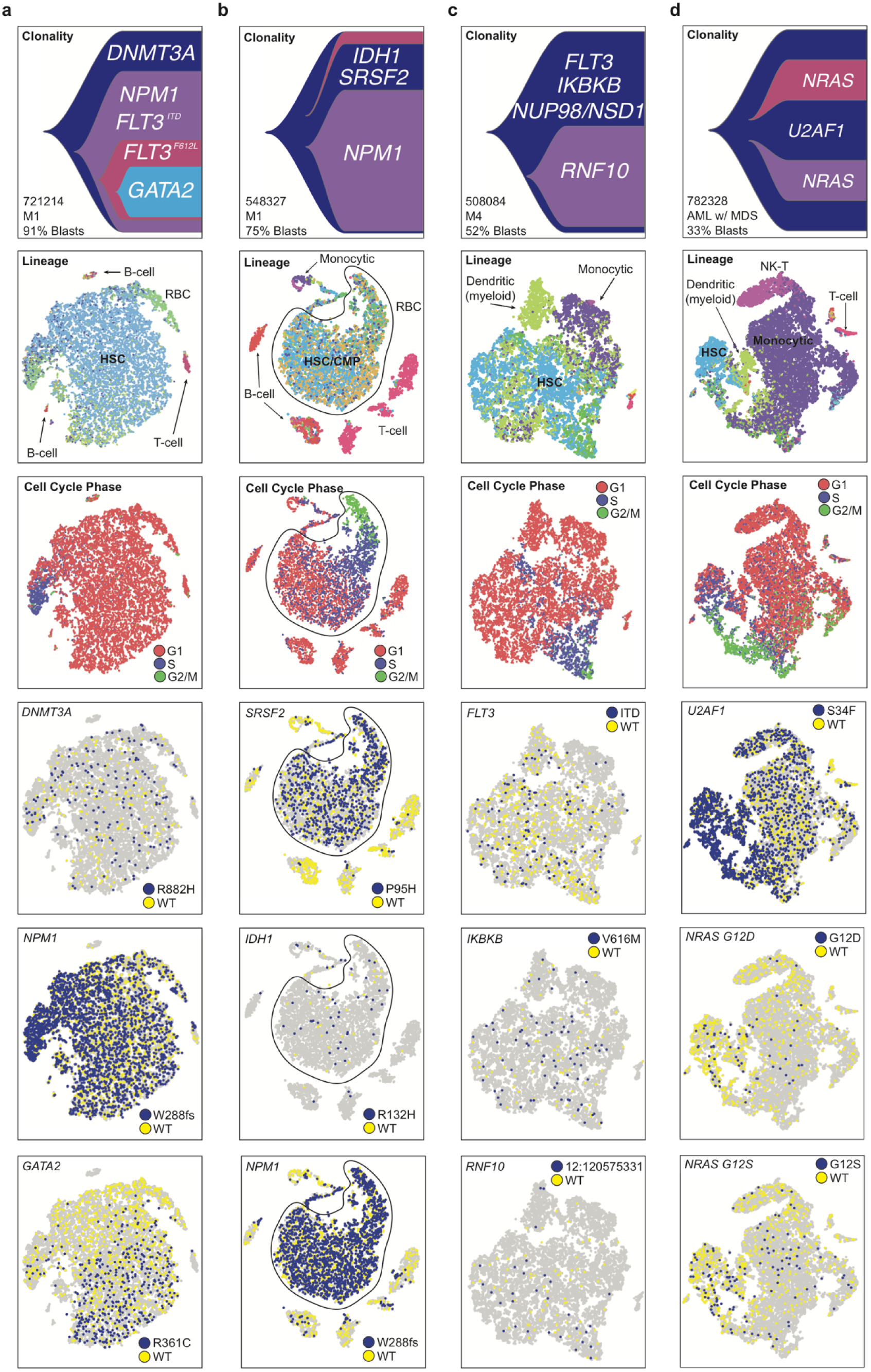
Single-cell mutation detection and interpretation in additional cases ordered by the differentiation signature of AML cells. (a) 721214, top to bottom: Clonality inferred from eWGS; cells colored according to cell cycle phase; cells colored according to single-cell genotype at the indicated site: blue, at least one mutant read detected; yellow, wild-type reads only; gray, no coverage. (b) 548327, putative AML cells circled. (c) 508084. (d) 782328.

We therefore combined single-cell mutation data with expression-based clustering and lineage inference to distinguish AML cells from non-AML clusters. Using the bone marrow sample from 809653 (which contained many non-AML cells, according to morphology and flow cytometry) we overlaid mutation data on the t-SNE plots by highlighting mutant cells (Fig. 2e-g). A germline SNP in the *BAG1* gene served as a positive control, marking SNP-containing cells in all expression clusters (Fig. 2h). By scRNA-seq, we detected cells expressing mutations in 8 genes, including *TP53*, *NRAS*, and *CEBPA* (Table 1, Supplementary Table 1). Several clusters were significantly enriched (*p* ≤ 0.05, one-sided Fisher exact test) for mutant cells; other cells in these clusters presumably contained undetected mutations (Fig. 2b-g). Two of these clusters were composed of cells that had stem/progenitor expression signatures (HSCs and MEPs). The other two were composed of cells expressing erythrocyte or monocyte markers; these cells would have been mistaken for normal cells using expression data alone. This was the only case with multiple copy number alterations, which provided additional sensitivity for defining AML cells (Supplementary Fig. 2c). In the other 4 cases, somatic mutations were also concentrated in specific cell clusters, suggesting that they represented AML cells (Fig. 3). This approach may miss small clusters with mutant cells, and rare AML cells that co-cluster with cells of different lineages. Overall, combining expression and mutation data delineated clusters of AML cells more comprehensively than either method alone, and allowed us to identify abnormally-differentiated AML cells (“lineage infidelity”).

By combining lineage inference with single-cell mutation identification, we estimated the extent of differentiation of each tumor. Our conclusions were supported by flow cytometry and morphology, but provided more insight into the differentiation state of AML cells in individual samples (Fig. 2, Fig. 3). In two cases (809653, 782328), a considerable fraction of the mutant cells had expression signatures consistent with differentiated cells: erythrocytes and monocytes in 809653 (Fig. 2c), and monocytes and NK-T cells in 782328 (Fig. 3d). Likewise, 548327 contained mutant cells that co-clustered with wild-type B- and T-cells, suggesting that some AML cells display lineage infidelity (Fig. 3b). Thus, this integrative genomic approach validates the concept that AML cells can have a variety of abnormal expression signatures, corresponding to different lineages and states of differentiation.

We next investigated the extent to which transcriptional heterogeneity was explained by mutational heterogeneity in each case. A subclonal mutation that drives an expression signature should be restricted in expression space. In contrast, a founding or subclonal mutation *not* associated with an expression signature should be present throughout expression space. Furthermore, this should not be dependent on restricted expression of the mutant gene. To this end, we highlighted mutant cells on the t-SNE plot of each sample, and identified mutations that are nonuniformly distributed, even after controlling for that gene’s expression (Fig. 2e-g, Fig. 3, Supplementary Fig. 7). The results reveal that the relationship between expression heterogeneity and mutational heterogeneity is case- and mutation-dependent.

Two cases, 721214 and 508084, contained subclonal mutations with nonuniform distributions (Fig 3a,c). Based on eWGS, 721214 contained a subclone defined by *GATA2*^R361C^. In the scRNA-seq data, cells expressing *GATA2*^R361C^ were predominantly restricted to one side of the t-SNE plot, suggesting that AML cells containing this mutation have a unique expression signature (Fig. 3a). We characterized this signature using multiple regression, and orthogonally confirmed its existence using flow cytometry and PCR (Methods, Supplementary Methods, Supplementary Discussion, Supplementary Fig. 8). Two cases (809653 (Fig. 2f-g) and 782328 (Fig. 3d)) exhibited complex mutation-associated expression profiles, and a third, 548327 (Fig. 3b), showed expression heterogeneity in the absence of discernable genetic heterogeneity (Supplementary Discussion).

The ability to map mutations in hundreds to thousands of individual cells also facilitates more conservative, direct analyses of intratumoral expression heterogeneity, using only cells that express a confirmed somatic mutation. We performed dimensionality reduction and graph-based clustering using these mutationally-defined AML cells, and selected genes with the most variable expression patterns (Supplementary Fig. 9). We averaged expression of these genes within each cluster, and hierarchically clustered the results (Supplementary Fig. 10). All samples showed intercellular heterogeneity in the expression of cell cycle genes (as expected) and genes that function in the immune system, especially the MHC Class II genes and/or *CD74*. All but one case (782328) showed intercellular variability in expression of *TP53*-interacting genes^15^. Three cases (508084, 548327, 721214) showed intercellular heterogeneity in genes that interact with the vascular cell adhesion gene *VCAM1*, and three (721214, 782328, 809653) showed heterogeneous expression of myeloid differentiation genes. There were also case-specific signatures, such as “response to reactive oxygen species” in 721214^15^. Notably, the *GATA2*^R361C^ expression signature is also evident in the mutant cells, suggesting an autoregulatory loop (Supplementary Discussion, Extended Data Fig).

Integrating approaches that link genetic and transcriptomic information in single cells has important implications for the study of heterogeneous cell populations. By combining eWGS and scRNA-seq data, we have shown that we can distinguish between tumor and non-tumor cells, identify tumor cells displaying lineage infidelity, more comprehensively evaluate the differentiation state of individual tumor samples, derive mutation-associated expression signatures, study transcriptional heterogeneity within confirmed tumor cells, and identify cell-surface markers that can be used to isolate specific cells for downstream studies. Further, the approach described here should be applicable--without additional modifications or customization--to virtually any tumor type.

## Methods

### Enhanced whole genome sequencing (eWGS), germline SNP detection, and somatic variant detection

For each case, we performed enhanced whole genome sequencing (eWGS) on bone marrow and matched normal tissue to identify germline and somatic variants. Libraries were captured using an IDT exome reagent enhanced with AML recurrently mutated genes^16^, then combined with WGS libraries and sequenced on an Illumina HiSeq4000, as described previously^17^. Germline mutations were called using GATK HaplotypeCaller v3.5^18^ (parameters - stand_emit_conf 10 -stand_call_conf 30) and filtered using recommended parameters (-- filterExpression "QD < 2.0 || FS > 60.0 || MQ < 40.0 || MQRankSum < −12.5 || ReadPosRankSum < −8.0"). Somatic mutations (SNVs, indels, and copy number alterations) were detected using an ensemble mutation calling approach, with detailed protocols as previously published^19^. Somatic structural variants were detected using Manta v0.29^20^.

### Bulk RNA-sequencing

RNA libraries were prepared using the TruSeq stranded kit, sequenced on the Illumina HiSeq platform, and aligned as described previously^19^. Expression quantification was performed using Kallisto 0.43.1 ^21^ and transcripts from ensembl version 74.

### Single-cell RNA-sequencing sample preparation, data generation, and coverage analysis

#### Flow sorting for live cells

Cryovials of AML cells were thawed as follows: While 9 ml of Fetal Bovine Serum (FBS) was allowed to come to ~24°C, AML cryovials were removed from liquid nitrogen, and warmed in a 37°C water bath until cells began to thaw. After 1 minute, 1 ml of room temperature FBS was added to the warming cryovial with a P1000 pipet tip and allowed to mix with thawing cells. The freshly added FBS was removed from the cell pellet and transferred back to the FBS stock. This process was repeated 3-4 times until all cells from the cryovial could be poured directly into the FBS stock. The empty cryovial was rinsed once more with the FBS mixture. Cells were then pelleted by centrifugation at 300 G for 5 minutes and resuspended in Phosphate Buffered Saline (PBS) at a concentration of 1×10^6^ cell/ml in 1x PBS. Cells were then pipetted through a 70 μm filter into a 5 ml tube for sorting. Cells were then stained with 1 μl 7-AAD per 1 ml of cells for 30 minutes at 4°C. If cell viability was ≤ 85%, stained cells were filtered through a 40 μM Flowmi cell strainer (Miltenyi), flow sorted, and gated using the FACS Chorus software (BD Biosciences).

#### 5-prime unbiased single-cell RNA library construction and sequencing

Cells were processed using the 10x Genomics Chromium controller and the Chromium Single Cell 5′ Library & Gel Bead Kit (PN 1000006) following the standard manufacturer’s protocols (https://tinyurl.com/y96l7lns). Two technical replicates were run in parallel for each sample. Briefly, between 14,000-21,000 live cells were loaded onto the Chromium controller in an effort to recover between 10,000-15,000 cells for library preparation and sequencing. Gel beads were prepared according to standard manufacturer’s protocols. Oil partitions of a single-cell + oligo coated gel beads (GEMs) were captured and reverse transcription was performed, resulting in cDNA tagged with a cell barcode and unique molecular index (UMI). Next, GEMs were broken and cDNA was amplified and quantified using an Agilent Bioanalyzer High Sensitivity chip (Agilent Technologies).

To prepare the final libraries, amplified cDNA was enzymatically fragmented, end-repaired, and polyA tagged. Fragments were then size selected using SPRIselect magnetic beads (Beckman Coulter). Next, Illumina sequencing adapters were ligated to the size-selected fragments and cleaned up using SPRIselect magnetic beads (Beckman Coulter). Finally, sample indices were selected and amplified for followed by a post sample index PCR double sided size selection using SPRIselect magnetic beads (Beckman Coulter). Final library quality was assessed using an Agilent Bioanalyzer High Sensitivity chip. Samples were then sequenced on the Illumina NovaSeq with a target of 150,000 reads/cell (2×150 paired end reads), yielding a median per-library depth of 192,427 reads per cell.

#### Evaluating transcript coverage as a function of distance from transcription start and stop points

Transcript alignment, counting, and inter-library normalization were performed using the Cell Ranger pipeline (10x Genomics, default settings, Version 2.1.1, GRCh38 reference) ^22^. For the genes *TP53*, *NPM1*, *GATA2*, and *DNMT3A,* the depth at each transcript was evaluated using both scRNA-seq data as well as bulk RNA-seq data. For each gene, a canonical isoform was chosen by consulting the APPRIS database^23^ (ENST00000445888.6, ENST00000296930.9, ENST00000341105.6 and ENST00000264709.7 respectively). For the scRNA-seq data, the number of unique barcode/UMI pairs was counted at each position. For the bulk RNA-seq data, the tool bamCoverage^24^ was used to generate a wiggle file over the transcript at 1bp bin size. The resulting tracks were visualized using the UCSC Genome Browser^25^. To reduce visual noise from intergenic reads, positions not overlapping the canonical isoform were not considered. Coverage plots for all mutated genes in this study are provided at https://github.com/genome/scrna_mutations.

To evaluate transcriptome-wide coverage, we used the annotation set GENCODE V27 to extract 20,090 genes with only one annotated isoform between 250bp and 11,000bp, with an average size of 1,569bp and median size of 829bp. Restricting to single isoform genes reduced noise related to alternative transcription start (TSS) and stop (TTS) sites. For each transcript in each sample in this study, single-cell transcriptome-wide coverage was quantified by counting the number of unique barcode/UMI pairs seen across the whole transcript. Then, for each position along the transcript, the number of unique pairs was divided by this total. This value was calculated as distance from the TSS for 5’ kit data, and distance from the TTS for 3’ kit data. To plot the results, the average value across all transcripts for all samples was calculated at each position. For shorter transcripts, positions with no data were not included in the average. The plot was also truncated to 10,000bp to avoid edge effects related to the transcript selection process. Coverage plots were generated using the Gviz^26^ and BiomaRt^27^ R packages, versions 1.22.3 and 2.34.2 respectively. For each locus, both coding and non-coding exonic nucleotides were considered at a 1bp bin size. Gene region tracks were retrieved directly from Ensembl v93. scRNA total read coverage was generated using bamCoverage, part of the deepTools package^24^, and scRNA cell barcode coverage can be found at https://github.com/genome/scrna_mutations.

#### Copy Number analysis

Gene expression matrices were analyzed with the CONICSmat package for R^28^. The default filtering and normalization procedures were followed, as outlined in https://github.com/diazlab/CONICS/wiki/Tutorial---CONICSmat;---Dataset:-SmartSeq2-scRNA-seq-of-Oligodendroglioma. The mixture model results were obtained, then restricted to regions of known copy number events from the eWGS with the best log-likelihood scores from the modelling: For sample 809653, these were chromosomes 1p and 7q. The z-scored posterior probabilities were clustered, using *k* = 4, and cell barcodes from the three clusters containing one or more of the expected events were gathered and visualized on the expression t-SNE plot (Supplementary Fig. 2c). High concordance was observed with expression-based classification of AML cells: 95.5% of cells classified as AML by copy number were also classified as AML by expression signature. (Conversely, 94.9% of cells classified as AML by expression were confirmed by CN).

#### Single-cell mutation identification and analysis

Using a Pysam-based tool (https://github.com/sridnona/cb_sniffer), we processed the aligned sequence data. For each cell barcode in the filtered Cell Ranger barcode list, and each somatic variant in the eWGS data, we identified all reads spanning the variant. Only cell-associated UMIs (defined as reads containing both a Chromium “Cellular Barcode” (CB) tag and a Chromium “Molecular Barcode” (UB) tag) were considered for downstream analysis. Variant positions were required to have a minimum base quality and mapping quality of at least 1. For each cell, we counted the number of unique reads matching the reference or variant allele. In rare cases where duplicate reads existed for a given UB and the base at the mutant position was not identical across all reads, we selected the most common base if it was present in at least 75% of the reads; otherwise all reads in the group corresponding to that UB were discarded. Several variants required additional steps to accurately identify mutant cells: Manual review revealed that two small indels in repetitive regions (*CEBPA* and *NPM1*) were frequently misaligned to several adjacent bases. This was resolved by parsing the bam cigar string to identify reads containing insertions or deletions at the appropriate locations using an additional pysam-based tool (https://github.com/genome/scrna_mutations/tree/master/misc_scripts). The large size of the characteristic large internal tandem duplication (ITD) in *FLT3* means that many reads containing the variant do not align correctly. We created a contig containing the variant sequence (+/- 250 bp), appended it to the reference, and realigned the scRNA data. Barcodes from reads uniquely aligning to the mutant *FLT3* sequence were then extracted. Similarly, the *NUP98*-*NSD1* fusion in 508084 was detected by appending the fusion transcript to the input GTF file, then using kallisto^21^ and its companion tool, pizzly, to identify fusion-supporting transcripts.

By assaying the positions of known somatic mutations in samples that did not harbor those mutations, we found that the false positive rate (the rate at which wild-type UMIs are called mutant) is site-specific, and at most 0.39% (Supplementary Table 4). For most SNVs, the vast majority of cells had at most one unique read at the variant position, but SNVs in several highly expressed, high-coverage genes (*U2AF1*, *NPM1*, *SRSF2*, *NRAS*) were more likely to have multiple reads per cell (Supplementary Fig. 2a). We then recorded which cells had wild-type or mutant sequence at that position. After using SciClone^29^ to assign each somatic variant to a subclone, we assigned mutation-containing cells (“mutant cells”) to their corresponding subclones. Cell-variant assignment can also be performed in an automated manner using the VarTrix tool (https://github.com/10xgenomics/vartrix).

#### Single-cell RNA-seq expression analysis and mutation integration

Transcript alignment, counting, and inter-library normalization were performed using the Cell Ranger pipeline (10x Genomics, default settings, Version 2.1.1). Using the Seurat R package^30^, cells that contained fewer than 10 expressed genes, more than 50% ribosomal transcripts, or more than 10% mitochondrial transcripts were removed. Genes that were expressed in fewer than 3 cells were also removed. For each cell, expression of each gene was normalized to the sequencing depth of the cell, scaled to a constant depth (10,000), and log-transformed. Variable genes were selected (x.low.cutoff = 0.0125, x.high.cutoff=5, y.cutoff=0.5, default settings otherwise). Principal component analysis was performed on the variable genes, and the optimal number of principal components (PCs) for each sample was chosen using a combination of elbow plots, jackstraw resampling, and PC expression heatmaps (508084: 6, 548327: 8, 721214: 5, 782328: 7, 809653: 6, 809653 AML cells: 6). PCs were used for dimensionality reduction if they explained at least 2% of the variance; were statistically significant according to jackstraw resampling; exhibited consistent expression variation in heatmaps; and were not composed entirely of ribosomal, mitochondrial, or immune genes. Dimensionality reduction and visualization were performed with the t-SNE algorithm (Seurat implementation) using the PCs selected above. Unsupervised graph-based clustering of cells was performed using the indicated PCs, with resolution = 0.7. Cell cycle phase was determined using methodology provided in Seurat, based on relative expression of phase-specific genes^3^. The distribution of mutations on the t-SNE plot was robust to filtering for mitochondrial and ribosomal transcripts, the number of PCs used, the clustering resolution, and normalization for cell cycle phase. The mutation distribution was also robust to the particular implementation of the t-SNE algorithm, with the Seurat and Cell Ranger implementations giving consistent results. To assess the relationship between mutation distribution and expression of the mutated gene, we colored each cluster in each t-SNE plot according to the expression-normalized mutant cell fraction (mutant cell fraction divided by the average expression of the mutant gene in that cluster).

Mutation-expressing cells were analyzed in isolation using analogous methods, with the exception that fewer PCs were required to capture the variability in the data (508084: 4, 548327: 3, 721214: 6, 782328: 7, 809653: 6).

#### Expression heatmaps

An expression heatmap was generated for each sample by selecting the top 10 genes in each of the top 20 PCs, and averaging the expression of each gene in each cluster. To connect heterogeneity to the graph-based clusters, and to examine relationships among clusters, we averaged expression within each cluster, and hierarchically clustered the results. For the analogous analysis performed on mutant cells in isolation, we used the top 20 genes from each of the top *n* PCs, where *n* was chosen separately for each sample to minimize noise (508084: 4, 548327: 3, 721214: 6, 782328: 7, 809653: 6).

#### Lineage inference and AML cell identification

Cell-type inference was performed in an unsupervised, marker-free manner by training a nearest-neighbor algorithm on expression data from the DMAP database^12^, using Spearman correlation as the distance metric. Using this approach, cells that co-cluster by graph-based clustering tend to have the same inferred lineage and express the corresponding cell-type markers (when known). In the case of AML cells, the assigned lineage represents the normal lineage to which the AML cell is most transcriptionally similar. To identify AML cells in highly heterogeneous samples (549327 and 809653), a one-sided Fisher exact test was used to identify cell clusters that were enriched for somatic mutations. In cases where most cells are AML cells, normal cell clusters were identified using a one-sided Fisher exact test for under-enrichment.

#### *GATA2*^R361C^ expression signature

Each cell containing a *GATA2*^R361C^ mutation was assigned to an expression cluster. Mutant cells were more highly concentrated in a contiguous group of expression clusters. To derive an expression signature for this mutation, we developed a regression model to identify genes whose expression varies with mutant cell concentration. For each gene *i*, multiple regression was used to quantify the relationship between mean expression (*E_i_*) and *GATA2*^R361C^ mutant cell fraction (*m*) across the 12 AML clusters, while controlling for mean cluster-wise *GATA2* expression (*g*): *E_i_* = *x_i_* + *y_i_m*+ *z_i_g*. After correcting for multiple hypotheses, we selected genes whose *p-*value for *y_i_* was at most 0.05.

#### Functional enrichment

Functional enrichment analyses were performed using ToppFun (https://toppgene.cchmc.org/enrichment.jsp) ^15^.

## Data Availability

Enhanced whole genome sequence (eWGS), bulk RNA-sequence, and single cell RNA-sequence (scRNA-seq) data generated during the current study are available in dbGaP (https://www.ncbi.nlm.nih.gov/gap/) with the primary accession code phs000159. The SRA IDs for this study are: SRR7904017, SRR7904018, SRR7904019, SRR7904020, SRR7910353, SRR7910351, SRR7910349, SRR7904016, SRR7903979, SRR7825447, SRR7825459, SRR7825446, SRR7825444, SRR7825491, SRR7825473, SRR7825453, SRR7825466, SRR7825499, SRR7825482, and SRR7939318.

## Author Contributions

Conceptualization: T.L., D.M.C.; Data Curation: A.P., S.W., C.M., I.F., S.S., D.M.C., T.L.; Data analysis and software: A.P., S.W., C.M., I.F., S.S.; Experimentation: D.Y.C., C.F., R.F.; Methodology: A.P., S.W., C.M., I.F., S.S., D.M.C., T.L.; Writing – original draft: A.P.; Writing – review and editing: A.P., S.W., C.M., I.F., S.S., D.M.C., T.L.; Funding acquisition and Supervision: D.M.C., T.L.

## Acknowledgements

Supported by the NCI K12 program (CA167540, R. Govindan, PI) to A. Petti, an R50 CA211782 to C. Miller, and an R35 CA197561 and P01 CA101937 to T. Ley. Matthew Christopher, MD, PhD and Amanda Smith, PhD developed the methods used for preparing cryovials of AML cells for analysis. We also thank Julia Lau for construction of the Chromium single cell gene expression libraries, and Jonathan Weinstein, PhD for comments on statistical methodology.

The authors declare the following competing interests: S.W., I.F., and D.M.C are employed by and hold shares in 10x Genomics.

Correspondence and requests for materials should be addressed to.

## Supplementary Figure Legends

**Supplementary Figure 1. Additional performance metrics for single-cell variant detection**.

(a) Distribution of variant-spanning reads for mutations in the indicated gene(s). (b) Log-scale distribution of variant-spanning reads for mutations in the indicated gene(s). (c) Relationship between single-cell and bulk RNA-seq VAF. (d) Mutant Cell Detection Rate as a function of gene expression in the single-cell data.

**Supplementary Figure 2. Clustering, overview of expression heterogeneity, and copy number analysis in 809653**. (a) t-SNE plot of scRNA-seq data, cells colored according to graph-based cluster assignment; putative AML clusters circled. (b) Hierarchical clustering of most heavily weighted genes in each principal component, averaged within graph-based clusters. (c) CNV analysis: blue, cells with detected CNVs; gray, no detected CNVs.

**Supplementary Figure 3. Clustering and overview of expression heterogeneity in 721214**.

(a) t-SNE plot of scRNA-seq data, cells colored according to graph-based cluster assignment; putative AML clusters circled. (b) Hierarchical clustering of most heavily weighted genes in each principal component, averaged within graph-based clusters.

**Supplementary Figure 4. Clustering and overview of expression heterogeneity in 548327**.

(a) t-SNE plot of scRNA-seq data, cells colored according to graph-based cluster assignment; putative AML clusters circled. (b) Hierarchical clustering of most heavily weighted genes in each principal component, averaged within graph-based clusters.

**Supplementary Figure 5. Clustering and overview of expression heterogeneity in 508084**.

(a) t-SNE plot of scRNA-seq data, cells colored according to graph-based cluster assignment; putative AML clusters circled. (b) Hierarchical clustering of most heavily weighted genes in each principal component, averaged within graph-based clusters.

**Supplementary Figure 6. Clustering and overview of expression heterogeneity in 548327**.

(a) t-SNE plot of scRNA-seq data, cells colored according to graph-based cluster assignment; putative AML clusters circled. (b) Hierarchical clustering of most heavily weighted genes in each principal component, averaged within graph-based clusters.

**Supplementary Figure 7. Clustered t-SNE plots colored according to expression-normalized mutation fraction in each cluster (selected genes).** (a) 809653, putative AML cells only. (b) 721214. (c) 548327. (d) 508084. (e) 782328.

**Supplementary Figure 8. *GATA2*^R361C^ Expression Signature in 721214.** (a) (left to right) t-SNE plot showing mutation-expressing cells in blue; cells colored according to graph-based cluster assignment; heatmap of mutation-dependent genes, with bar graph showing mutant cell fraction in each cluster. (b) Cells colored according to *VIM* expression, and scatterplot showing average *VIM* expression in each cluster as a function of the mutation fraction of each cluster. (c) t-SNE plot constructed from mutant cells, which are colored according to the mutation they contain: *GATA2*^R361C^, yellow; *DNMT3A*^R882H^, pink; *FLT3*-ITD, green; *FLT3*^F612L^, purple; *NPM1*^W288FS^; other somatic mutation, gray. (d) Cells colored according to *CD99* expression, and scatterplot showing average *VIM* expression in each cluster as a function of the mutation fraction of each cluster. (e) Gating of cells based on *CD99* expression using flow cytometry. (f) Variant allele fraction of the founding clone *DNMT3A* R882H mutation and the subclonal *GATA2* R361C mutation in unsorted cells (gray), CD99-high cells (blue), and CD99-low cells (red).

**Supplementary Figure 9. Dimensionality reduction, clustering, and lineage inference of mutant cells.** (a) 809653. (b) 721214. (c) 548327. (d) 508084. (e) 782328.

**Supplementary Figure 10. Cluster-averaged gene expression profiles of variable genes in mutant cells.** (a) 809653. (b) 721214. (c) 548327. (d) 508084. (e) 782328.

